# Disease context dictates the cellular targets of IL-17 in inflammatory skin disease

**DOI:** 10.64898/2026.03.23.713409

**Authors:** Kellen J. Cavagnero, Fengwu Li, Haley Jo, Carlos Aguilera, Jennifer Fox, Joseph Kirma, Rachael Bogel, J. Michelle Kahlenberg, Lam C. Tsoi, Johann E. Gudjonsson, Richard L. Gallo

## Abstract

Therapeutic blockade of IL-17 and TNF can effectively treat inflammatory skin diseases such as hidradenitis suppurativa and psoriasis, yet the relative importance of the different cell types that respond to IL-17 and TNF remains unresolved. Keratinocytes are viewed as the dominant effector cells, whereas fibroblasts have recently emerged as important contributors. In mice, topical imiquimod induces IL-17- and TNF-dependent skin inflammation and is frequently used to model psoriasis. Here, we demonstrate that intradermal injection of recombinant IL-17 and TNF elicits skin inflammation with features of hidradenitis suppurativa, including a gene expression program that is distinct from psoriasis and imiquimod-induced inflammation. Single-cell transcriptomic network analysis identified dermal fibroblasts as the dominant cell communication hub in hidradenitis suppurativa and in mice injected with IL-17 and TNF. In contrast, fibroblasts and keratinocytes both show strong network involvement in psoriasis and in mice challenged with imiquimod. Cell-type-specific deletion of IL-17 receptor A in mice revealed that imiquimod-induced inflammation depends equally on IL-17 signaling in fibroblasts and keratinocytes, whereas inflammation induced by intradermal IL-17 and TNF only requires fibroblasts to recognize IL-17 and is independent of keratinocyte IL-17 sensing. Single-cell transcriptomic analysis of these conditional knockout mice further demonstrated that keratinocytes and fibroblasts activate divergent and disease-dependent transcriptional programs following activation by IL-17. Together, these findings introduce a new conceptual framework wherein IL-17 signaling is routed through distinct cellular and molecular pathways depending on disease context and establish complementary experimental systems for interrogating type 17 skin inflammation.

## Introduction

Psoriasis and hidradenitis suppurativa (HS) are chronic inflammatory skin diseases that have strikingly different clinical and histological patterns but are both characterized by type 17 immune responses. In psoriasis vulgaris, inflammation is relatively mild, largely confined to the epidermis, and comprises neutrophils, lymphocytes, dendritic cells, and macrophages^1^. In HS, an intense cellular infiltrate of neutrophils, lymphocytes (including B/plasma cells), dendritic cells, and macrophages is present and localized to the deep dermis^2^. Despite these differences, therapeutic blockade of IL-17 and TNF can effectively treat both diseases, raising a central unresolved question: How can the same inflammatory cytokines drive fundamentally distinct patterns of tissue inflammation?

Current conceptual models of type 17 skin inflammation position keratinocytes (KCs) as the dominant tissue-resident target of IL-17 and TNF. IL-17 and TNF act on KCs to promote epidermal hyperplasia, chemokine production, and neutrophil recruitment, establishing a feed-forward inflammatory loop at the skin surface^3, 4^. These models have been shaped in part by studies of the topical imiquimod (IMQ) mouse model, which induces IL-17- and TNF-dependent epidermal inflammation characteristic of psoriasis^5^. However, no experimental system has been shown to reproduce the robust dermal type 17 inflammation observed in HS, limiting our ability to interrogate the pathophysiological mechanisms of disease contexts dominated by dermal immune responses. Instead, findings from the IMQ model have been extrapolated to HS^6^, despite the different inflammatory characteristics.

Dermal fibroblasts (FBs) are highly abundant tissue-resident cells. Long regarded primarily as structural cells, FBs have recently emerged as active participants in immune responses^7^. Single-cell transcriptomic analyses have identified subsets of immune-acting FBs (IAFs) that express cytokines, chemokines, and antimicrobial genes^8, 9, 10^. *In vitro*, FBs can recognize IL-17 and TNF, secrete chemokines, and recruit neutrophils^8, 9, 10^. *In vivo* studies using conditional knockout mice have demonstrated that FB signaling through IL-17RA and TNFR1 contributes to neutrophil recruitment during IMQ-induced skin inflammation^8, 9^. Analyses of human psoriasis skin before and after therapeutic cytokine blockade further supported a role for FBs as important targets of IL-17 and TNF signaling^8, 9, 11^.

Together, these findings show that both KCs and FBs promote type 17 skin inflammation; however, their relative contributions—and whether these contributions vary across disease contexts, remains unresolved. In particular, it is unknown whether IL-17 and TNF are routed through distinct tissue-resident cell types depending on whether inflammation is epidermal or dermal.

Here, we address this question by integrating human transcriptomics with murine experimental systems. We find that intradermal (i.d.) delivery of IL-17 and TNF elicits a dermal inflammatory program that resembles HS inflammation and is distinct from IMQ-induced and psoriasis inflammation. Using cell-type-specific genetic perturbations and machine learning network analysis, we demonstrate that FBs play a dominant role in dermal inflammation, whereas FBs and KCs have similar contributions in epidermal inflammation. These findings reveal that disease context dictates the cellular pathways through which IL-17 drives skin inflammation and establish complementary experimental systems for dissecting disease-specific type 17 inflammatory mechanisms.

## Results and Discussion

### Establishment of experimental systems of epidermal versus dermal type 17 skin inflammation in mice

To study how IL-17 and TNF drive different patterns of skin inflammation, we first established murine experimental systems that reflect the different inflammatory patterns observed in psoriasis and HS. Topical application of IMQ to mouse dorsal (back) skin represents a well-described model of psoriasis, inducing IL-17- and TNF-dependent epidermal inflammation^5^. In contrast, we hypothesized that i.d. delivery of recombinant IL-17A and TNF (IL-17/TNF) would elicit robust inflammation that is restricted to the deep dermis and more characteristic of HS (Fig. 1A).

**Fig. 1.**
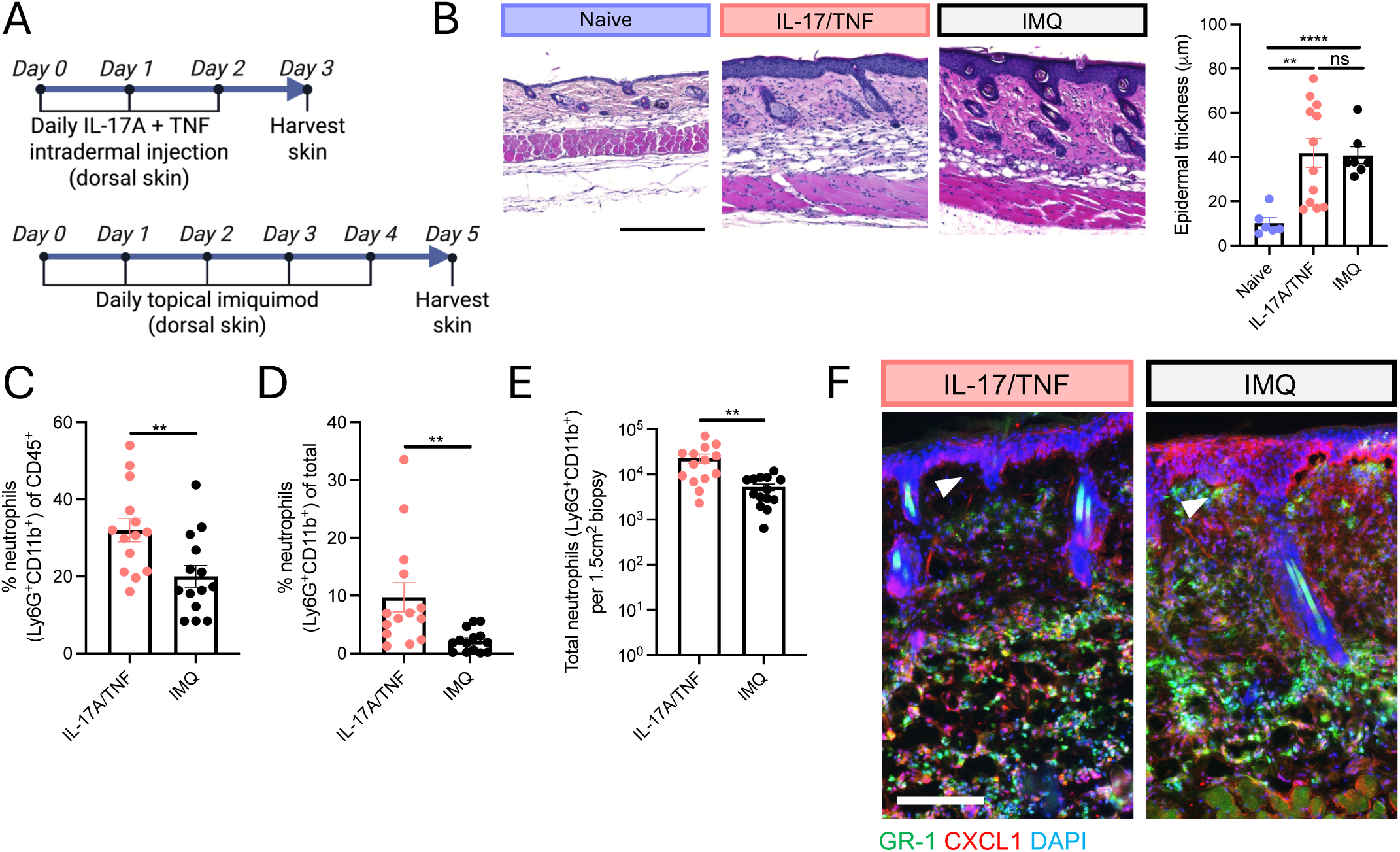
Establishment of experimental systems of epidermal versus dermal type 17 skin inflammation in mice. (A) Murine type 17 skin inflammation experimental systems. (B) Representative back skin H&E staining (left) and epidermal thickness quantification (right). Flow cytometric quantification of the frequency of neutrophils (Ly6G+CD11b+) of live CD45+ skin cells (C), frequency of neutrophils of total live skin cells (D), and total neutrophils per 1.5cm2 skin biopsy (E). (F) Skin immunostaining. Arrows denote region with marked difference in sub-epidermal inflammation. Scale bar, 150um. Data are represented as mean +/- SEM. ns (not significant), **p < 0.01, and ****P < 0.0001 using unpaired t-test. IMQ, imiquimod.

Histological analysis confirmed that both IMQ and IL-17/TNF caused skin inflammation and epidermal hyperplasia (Fig. 1B). Flow cytometric analysis identified neutrophil accumulation in both contexts, with substantially greater neutrophilia following i.d. IL-17/TNF delivery (Fig. 1C-E). Notably, staining for GR-1 (high expressed by neutrophils and weakly by monocytes and macrophages^12^) localized predominantly to the reticular (deep) dermis following IL-17/TNF injection and was absent from the papillary (upper) dermis, whereas IMQ-induced GR-1+ cells were positioned proximal to the epidermis (Fig. 1F), mirroring the distinct spatial distribution of inflammatory infiltrate in HS versus psoriasis. The neutrophil chemokine CXCL1 was highly expressed throughout the skin layers in response to IMQ but was restricted to the deep dermis following IL-17/TNF injection (Fig. 1F). These epidermal and dermal patterns of neutrophil chemokine expression and inflammatory infiltrate were also observed with topical and i.d. microbial triggers (Fig. S1A-B).

### Distinct cellular and molecular features underlie dermal and epidermal inflammation in mouse and human

To identify the tissue-resident cell populations coordinating these distinct patterns of type 17 skin inflammation, we performed scRNA-seq followed by unbiased cell-cell communication analysis with CellChat^13^. In the murine dermal inflammatory context, FBs emerged as the dominant signaling hub, whereas both FBs and KCs showed strong network activity in the epidermal inflammatory context (Fig. 2A), consistent with CXCL1 immunostaining (Fig. 1F). Notably, KC-FB interactions were prominent in the IMQ context, suggesting a role for epidermal-dermal crosstalk in type 17 inflammation.

**Fig. 2.**
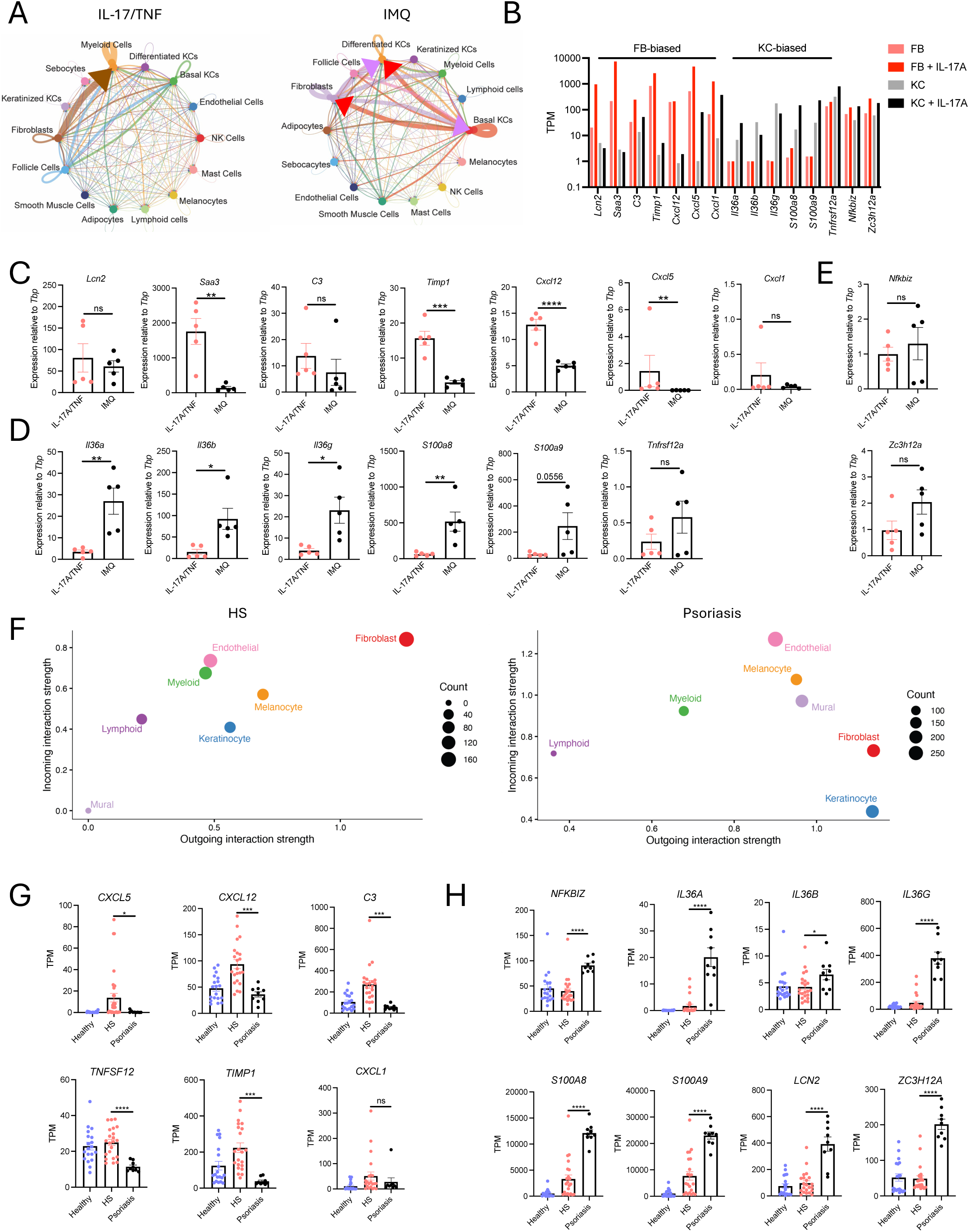
Distinct cellular and molecular features underlie dermal and epidermal inflammation in mouse and human. (A) scRNA-seq cell-cell communication analysis of mouse back skin challenged with intradermal IL-17/TNF (left) and topical IMQ (right) using CellChat. (B) RNA-seq gene expression of select IL-17 responsive genes in primary mouse FB and KC treated with IL-17A. qPCR gene expression of select IL-17 response genes that are FB-biased (Fig. 2C), KC-biased, (Fig. 2D) and non-biased (Fig. 2E) in mouse back skin challenged with IL-17/TNF versus IMQ. (F) scRNA-seq cell-cell communication analysis of skin from human HS (left) and psoriasis (right). RNA-seq gene expression of select IL-17 responsive genes enriched in skin from human HS (G) versus psoriasis (H). Data are represented as mean +/-SEM. ns (not significant), *p < 0.05, **p < 0.01, ***P < 0.001, and ****P < 0.0001 using unpaired t-test. Fibroblast, FB; keratinocyte, KC; IMQ, imiquimod; HS, hidradenitis suppurativa; TPM, transcript per million.

We next sought to understand how IL-17 directly affects FBs relative to KCs. To this end, we performed RNA-seq on primary mouse dermal FBs cultured with IL-17A and compared this response to that of primary mouse epidermal KCs cultured with IL-17A (GSE129176^14^). This analysis revealed three modules of key IL-17-responsive genes: one highly expressed by FBs, one highly expressed by KCs, and one highly expressed by both cell types (Fig. 2B). Specifically, FBs activated by IL-17 expressed markedly greater levels of *Lcn2*, *Cxcl12*, *Cxcl5*, *C3*, and *Timp1*, whereas KCs activated by IL-17 expressed substantially greater levels of *Il36a*/*b*/*g*, *S100a8*/*9*, and *Tnfrsf12a* (encodes LIGHT). Baseline expression and induction of *Nfkbiz* and *Zc3h12a* (encodes Regnase-1) was similar between the two cell types.

Transcriptomics of mouse back skin challenged with topical IMQ versus i.d. IL-17/TNF was next performed to begin understanding the relative contribution of FB and KC recognition of IL-17 *in vivo*. Investigation of IL-17 response genes that are FB-biased (Fig. 2C), KC-biased, (Fig. 2D) and non-biased (Fig. 2E) revealed that many key type 17 inflammatory genes were differentially expressed between the two experimental systems. Genes upregulated during dermal inflammation included *Saa3*, *Timp1*, *Cxcl5*, and *Cxcl12*, whereas genes upregulated during epidermal inflammation included *Il36a/b/g* and *S100a8/9*. Thus, the dermal type 17 inflammation transcriptome is consistent with relatively more FB activation by IL-17, whereas the epidermal type 17 inflammation transcriptome is consistent with more KC activation by IL-17.

To understand whether the cellular and molecular features underlying these experimental systems translate to human type 17 skin inflammation, we next performed a comparative analysis of HS and psoriasis skin transcriptomes. We first analyzed scRNA-seq datasets from HS (GSE175990^15^) and psoriasis (GSE162183^16^) skin and performed unbiased cell-cell communication analysis using CellChat^13^. FBs dominated the communication network in HS skin, while psoriasis skin exhibited strong contributions from both FBs and KCs (Fig. 2F). These results were analogous to those in murine dermal and epidermal type 17 inflammation experimental systems, respectively. Analysis of bulk RNA-seq data from lesional HS (GSE154773^17^) and psoriasis (GSE205748^18, 19^) skin revealed that the mouse FB/dermal-associated IL-17 response genes were largely upregulated in HS skin compared to psoriasis (Fig. 2G), whereas the mouse KC/epidermal-associated IL-17 response genes were enriched in psoriasis skin (Fig. 2H).

Together, these data establish i.d. IL-17/TNF delivery in mice as an experimental system with several histological, cellular, and molecular features corresponding to human dermal type 17 inflammation in HS, while the topical IMQ system more closely resembles human epidermal type 17 inflammation in psoriasis. Notably, the IL-17/TNF system does not recapitulate aspects of HS beyond type 17 inflammation, including formation of epithelial tunnels and tertiary lymphoid structures (TLS). The distinguishing features of the IMQ and IL-17/TNF systems led us to hypothesize that FB recognition of IL-17 is more important for dermal type 17 inflammation, whereas KC recognition of IL-17 is more important for epidermal type 17 inflammation.

### IL-17 signaling in fibroblasts and keratinocytes is equally required for epidermal type 17 inflammation

To directly assess the relative requirement of FB and KC recognition of IL-17 in dermal and epidermal type 17 inflammation, we used the Cre-Lox strategy to generate conditional knockout mice lacking IL-17 receptor A (IL-17RA) selectively in FBs (*Pdgfra*^Δ^*^Il17ra^*) or KCs (*Krt14*^Δ^*^Il17ra^*). This IL-17 receptor subunit is required for the activity of all IL-17 family members involved in type 17 inflammation (IL-17A/F/C/D). *Pdgfra* and *Krt14* Cre drivers have been extensively studied and are highly specific to FB and KC lineages, respectively^8, 10, 20, 21, 22, 23^.

We first subjected *Pdgfra*^Δ^*^Il17ra^*, *Krt14*^Δ^*^Il17ra^*, and Cre^-^ littermate control mice to topical IMQ application (Fig. 3A). Consistent with prior literature^3^, removal of IL-17RA in KCs reduced epidermal thickness by histological analysis (Fig. 3B). Removal of IL-17RA in FBs also significantly reduced epidermal thickness and did so to the same extent as removal of IL-17RA in KCs. This epidermal alteration following FB perturbation demonstrates functional FB-to-KC communication. Flow cytometric analysis indicated that the frequency and total number of neutrophils in the skin were reduced to the same extent in both IL-17RA conditional knockout strains compared to controls (Fig. 3C-E). Ly6G immunostaining validated the attenuation of skin neutrophilia in *Pdgfra*^Δ^*^Il17ra^* and *Krt14*^Δ^*^Il17ra^* mice (Fig. 3F).

**Fig. 3.**
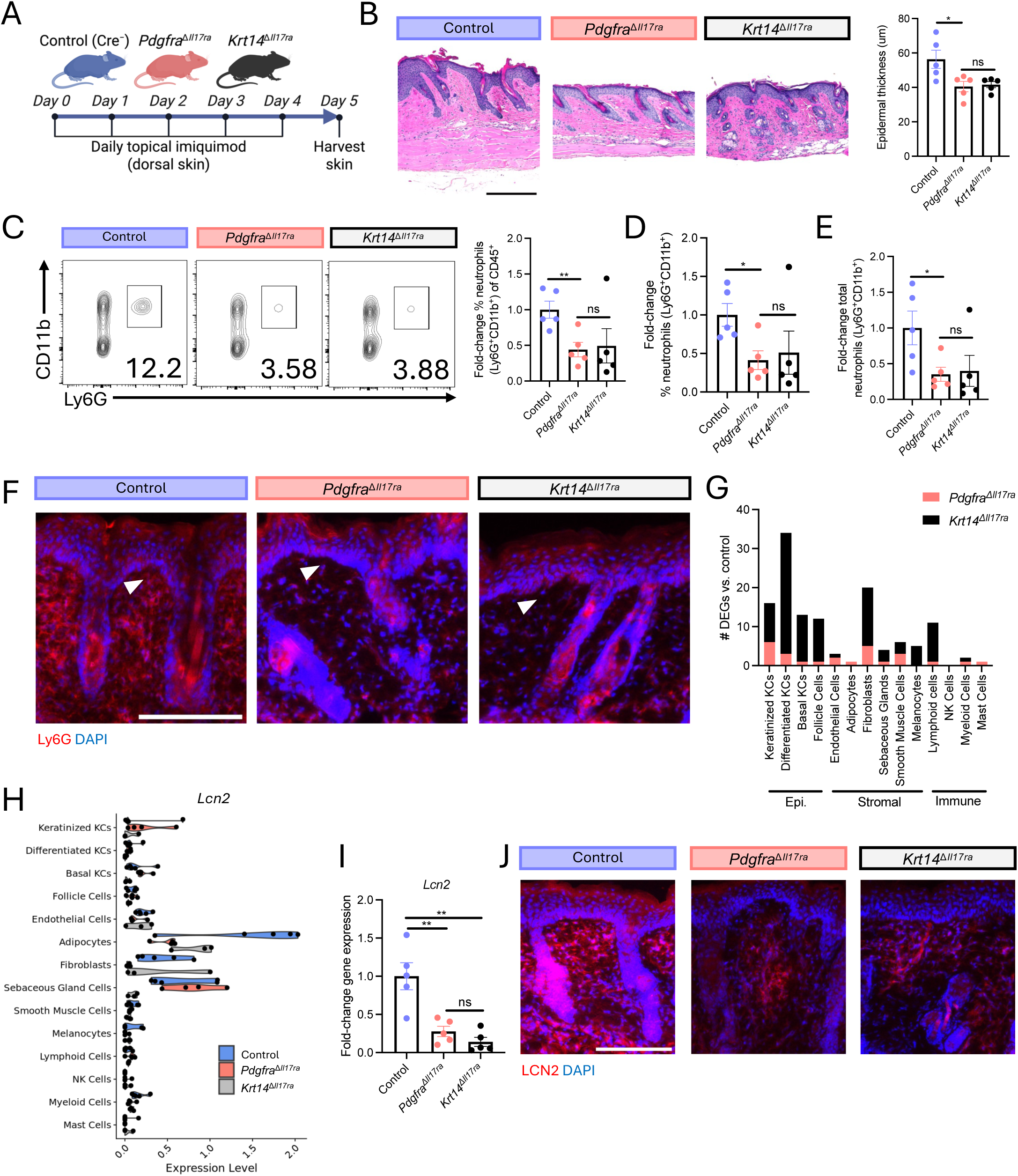
IL-17 signaling in fibroblasts and keratinocytes is equally required for epidermal type 17 inflammation. (A-J) Topical IMQ experimental system. Control = Cre^-^ littermates. (A) Experimental schematic. (B) Representative H&E staining (left) and epidermal thickness quantification (right). Flow cytometric quantification of the fold-change frequency of neutrophils (Ly6G+CD11b+) of live CD45+ skin cells (C), frequency of neutrophils of total live skin cells (D), and total neutrophils per 1.5cm2 skin biopsy (E). (F) Representative skin immunostaining. Arrows denote region with marked difference in sub-epidermal inflammation. Scale bar, 150um. (G) scRNA-seq pseudo-bulk DEGs between condition knockout mice and Cre^-^ control skin (P_adj_<0.05). N=4-6mice/group. (H) scRNA-seq pseudo-bulk gene expression. Each point represents 1 mouse. (I) Skin qPCR gene expression. (J) Representative skin immunostaining. Scale bar, um. ns (not significant), *p < 0.05, and **P < 0.01 using unpaired t-test. Differentially expressed genes, DEGs; IMQ, imiquimod. Epi, epidermal; KC, keratinocyte; NK, natural killer.

To probe the molecular basis of these findings, we performed scRNA-seq of the skin from *Pdgfra*^Δ^*^Il17ra^*, *Krt14*^Δ^*^Il17ra^*, and Cre^-^ control mice following IMQ challenge. Pseudo-bulk differential expression analysis demonstrated that the transcriptional response to IMQ depended on IL-17 signaling in both KCs and FBs (Fig. 3G). However, the composition, magnitude, and cellular distribution of the transcriptional changes differed markedly depending on which cell type lacked IL-17 signaling. Notably, disruption of IL-17 signaling in either compartment altered the transcriptional program of the other, further supporting functional crosstalk between FBs and KCs. Deletion of IL-17RA in KCs produced the strongest effect, predominantly altering KC gene expression and secondarily affecting FBs. This perturbation reduced KC expression of a subset of the IL-17-responsive genes identified *in vitro*, including *S100a8/9*, *Zc3h12a*, and *Il36* (Fig. S2A). It also suppressed FB expression *Lcn2* (Fig. 3H)— a neutrophil chemokine^24, 25^ that is required for cutaneous neutrophilia^25, 26, 27, 28^, upregulated at the protein level in psoriasis and HS skin^25, 29^, and highly IL-17-responsive in FBs in culture. In contrast, deletion of IL-17RA in FBs led to more balanced transcriptional changes across FBs and KCs. FB-specific deletion of IL-17RA strongly reduced expression of a small subset of FB IL-17-responsive genes identified *in vitro*, including *Lcn2*, *Saa3, and Timp1*, while exerting comparatively modest effects on KC-biased IL-17 target genes (Fig. 3H, S2A). These changes in gene expression were validated by qPCR (Fig. 3I, S2B). Consistent with transcriptional findings, immunostaining confirmed that loss of IL-17 signaling in KC or FB reduced dermal LCN2 protein expression (Fig. 3J).

Skin from different anatomical regions has different proportions of KCs-to-FBs. We hypothesized that the role of KCs may be more pronounced in skin from a region with a relatively high KC-to-FB ratio, such as mouse ear skin. To test this, we subjected *Pdgfra*^Δ^*^Il17ra^*, *Krt14*^Δ^*^Il17ra^*, and Cre^-^ control mouse ear skin to topical IMQ application. Removal of IL-17RA in FBs and KCs reduced skin inflammation, epidermal thickness, and *Lcn2* expression (Fig. S2C-D). No significant difference was observed between *Pdgfra*^Δ^*^Il17ra^* and *Krt14*^Δ^*^Il17ra^* mice, demonstrating equal requirement for KC and FB recognition of IL-17.

Taken together, these results demonstrate that IMQ-induced IL-17 recognition by KCs and FBs drives divergent transcriptional responses that are similarly required for skin neutrophilia and epidermal hyperplasia, regardless of the anatomical region. Prior studies comparing mice with global deletion of IL-17RA in mice to strains lacking IL-17RA specifically in KCs, T cells, neutrophils, and macrophages in the IMQ context concluded that KCs are the dominant target of IL-17 in the skin as only *Krt14*^Δ^*^Il17ra^* mice phenocopied global knockout animals^3^. This study reinforced conceptual models of psoriasis that emphasize KCs as the principal targets of IL-17 signaling^30^. However, this study and others did not investigate the relative contribution of FBs. Through direct and comprehensive comparison of *Pdgfra*^Δ^*^Il17ra^* and *Krt14*^Δ^*^Il17ra^* mice, our findings refine the model of epidermal type 17 inflammation to recognize equal contribution of KC and FB recognition of IL-17 and bi-directional KC-FB communication.

### Dermal type 17 inflammation requires IL-17 signaling in fibroblasts but not keratinocytes

We next subjected *Pdgfra*^Δ^*^Il17ra^*, *Krt14*^Δ^*^Il17ra^*, and Cre^-^ littermate control mice to i.d. IL-17/TNF injection (Fig. 4A). In contrast to the IMQ-induced epidermal inflammatory context, removal of IL-17RA in KCs had no detectable effect on epidermal morphology but removal of IL-17RA in FBs significantly reduced epidermal thickening (Fig. 4B). Flow cytometric analysis demonstrated cutaneous neutrophilia was significantly reduced in *Pdgfra*^Δ^*^Il17ra^* mice compared to Cre^-^ controls, whereas *Krt14*^Δ^*^Il17ra^* mice had similar neutrophil levels compared to Cre^-^ controls (Fig. 4C-E). Ly6G immunostaining confirmed that neutrophilia was reduced in *Pdgfra*^Δ^*^Il17ra^* but not *Krt14*^Δ^*^Il17ra^* mice (Fig. 4F).

**Fig. 4.**
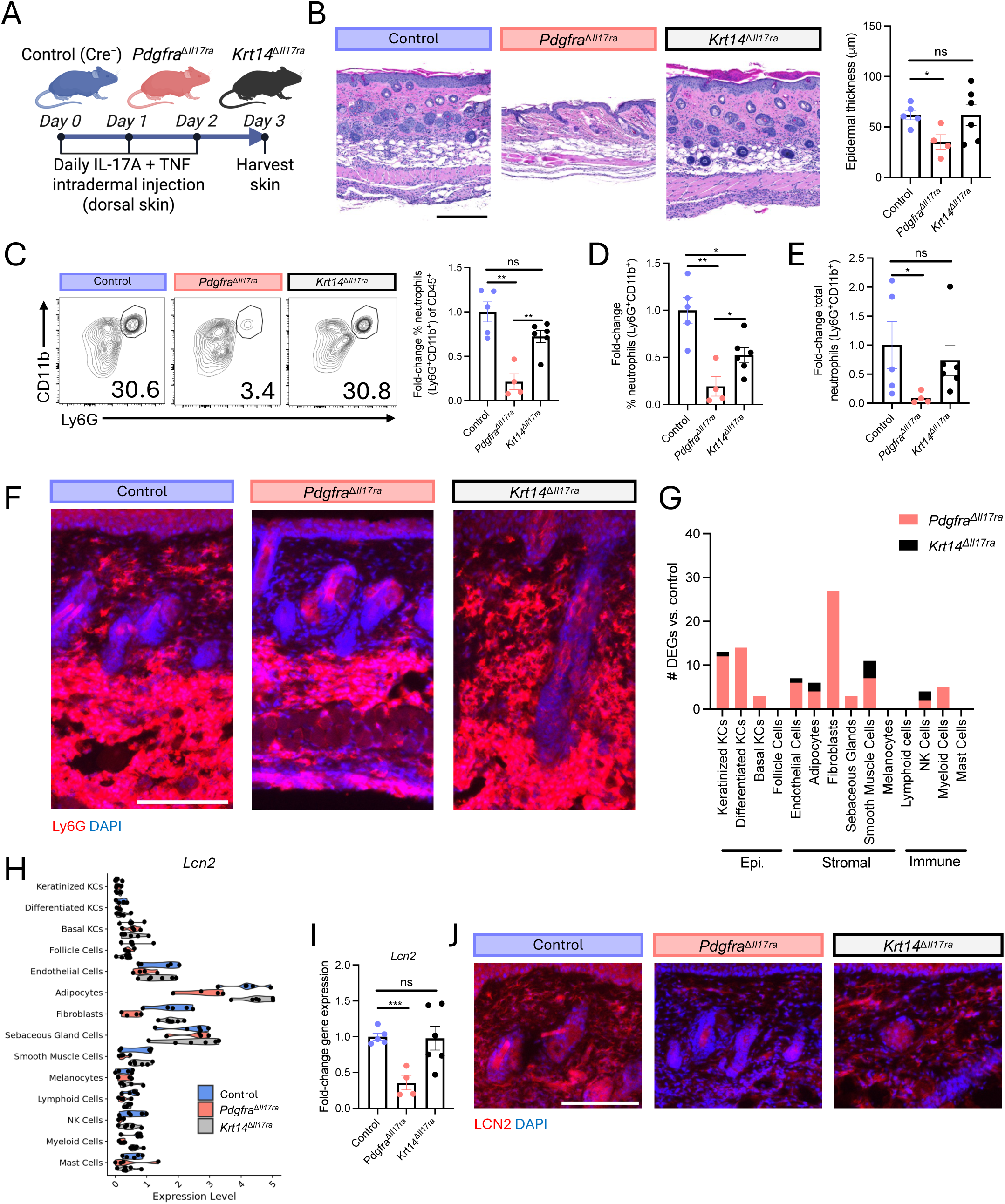
Dermal type 17 inflammation requires IL-17 signaling in fibroblasts but not keratinocytes. (A-J) i.d. IL-17/TNF experimental system. Control = Cre^-^ littermates. (A) Experimental schematic. (B) Representative H&E staining (left) and epidermal thickness quantification (right). Flow cytometric quantification of the fold-change frequency of neutrophils (Ly6G+CD11b+) of live CD45+ skin cells (C), frequency of neutrophils of total live skin cells (D), and total neutrophils per 1.5cm2 skin biopsy (E). (F) Representative skin immunostaining. Scale bar, 150um. (G) scRNA-seq pseudo-bulk DEGs between condition knockout mice and Cre^-^ control skin (P_adj_<0.05). N=4-6mice/group. (H) scRNA-seq pseudo-bulk gene expression. Each point represents 1 mouse. (I) Skin qPCR gene expression. (J) Representative skin immunostaining. Scale bar, um. ns (not significant), *p < 0.05, **p < 0.01, and ***P < 0.001 using unpaired t-test. Differentially expressed genes, DEGs; i.d., intradermal; Epi, epidermal; KC, keratinocyte; NK, natural killer.

To define the molecular basis of these findings, we performed scRNA-seq on *Pdgfra*^Δ^*^Il17ra^*, *Krt14*^Δ^*^Il17ra^*, and Cre^-^ mouse skin following IL-17/TNF challenge. Pseudo-bulk differential expression analysis demonstrated that the transcriptional response to IL-17/TNF depended overwhelmingly on IL-17 recognition by FBs, not KCs (Fig. 4G). Deletion of IL-17RA in FBs reduced FB expression of many of the IL-17-responsive FB genes identified *in vitro*, including *Lcn2*, *Cxcl12*, *Cxcl5*, *Saa3,* and *Timp1* (Fig. 4H, S2E). This reduction was validated by qPCR (Fig. 4I, S2F). Notably, disruption of IL-17 signaling in FBs altered the transcriptional program of the epidermal compartment, supporting functional FB-to-KC communication. Consistent with transcriptional findings, immunostaining confirmed that loss of IL-17 signaling in FBs reduced dermal LCN2 protein expression (Fig. 4J).

Histologic analysis revealed neutrophil accumulation within the adipose compartment of the deep dermis following i.d. IL-17/TNF administration (Fig. 4F), and molecular analysis of dermal type 17 inflammation identified robust expression of several inflammatory genes in adipocytes, including *Lcn2*, *Saa3*, and *Cxcl12* (Fig. 4H, S2E). Subsets of FBs represent the dominant *in vivo* adipocyte progenitor^31^. Therefore, adipocytes may contribute to the phenotype observed in *Pdgfra*^Δ^*^Il17ra^* mice. To address this possibility, we first performed experiments with cultured primary mouse dermal FBs and FB-derived adipocytes (Fig. S3A). RNA-seq demonstrated that both cell states highly expressed neutrophil chemokines in response to IL-17/TNF (Fig. S3B). Consistently, both FB and adipocytes were activated by IL-17/TNF to recruit neutrophils in a transwell migration assay (Fig. S3C). We next generated mice lacking IL-17RA specifically in adipocytes using *Adipoq*-Cre^32^, which is expressed at later stages of adipocyte differentiation and distinguishes mature adipocytes from FBs. Compared to Cre^-^ controls, *Il17ra* expression in *Adipoq*^Δ^*^Il17ra^* mice was significantly reduced in adipose tissue but unchanged in isolated epidermis and dermal FBs (Fig. S3D). We next challenged *Adipoq*^Δ^*^Il17ra^* and Cre^-^ controls with i.d. IL-17/TNF. While *Adipoq*^Δ^*^Il17ra^* mice demonstrated reduced *Lcn2* expression compared to Cre^-^ controls (Fig. S3E), no significant decrease was observed in other key IL-17 response genes, cutaneous neutrophilia, epidermal hyperplasia (Fig. S3F-I). These findings indicate that adipocyte recognition of IL-17 is dispensable for dermal type 17 inflammation.

Together, these results demonstrate that FB recognition of IL-17 is critical for dermal type 17 inflammation, including secondary epidermal hyperplasia, whereas IL-17 signaling in KCs and adipocytes is dispensable. Thus, type 17 skin inflammation can exist independent of direct KC IL-17 sensing, positioning FBs as a critical driver of dermal type 17 inflammation. This conclusion is consistent with recent evidence revealing an important role for FBs in HS. In 2024, three independent studies identified subsets of immune-acting FBs enriched in HS^33,34,35^. While these studies focused on the role of immune-acting FBs in TLS formation, one study noted that these cells are a dominant source of neutrophil chemokine^34^. Further, a 2025 study identified that *NCSTN*, a component of the γ-secretase complex linked to monogenic HS, is selectively downregulated in dermal FBs of sporadic HS patients, and that its loss potentiates FB—but not KC—neutrophil chemokine response to TNF^36^.

Collectively, our comprehensive analysis of human and mouse epidermal and dermal type 17 inflammation supports a revised framework of type 17 inflammation in which IL-17 signaling is not routed through a fixed effector cell type but is instead dictated by disease context. Dermal type 17 inflammation is driven by FB IL-17 recognition, whereas epidermal type 17 inflammation requires coordinated IL-17 signaling in FBs and KCs. This work suggests KCs are not a major target of IL-17 in dermal type 17 inflammation as in HS. The context-dependence of KC and FB response to IL-17 in human disease may be explained by functional differences in upstream IL-17 producing cells that has been described in HS and psoriasis^37^. Dermal-predominant immune infiltrates are observed in many human skin diseases, including neutrophilic dermatosis^9^. Therefore, the FB-centered paradigm we describe may represent a general principle for deep-tissue inflammation, and the experimental systems we have established may be useful for further mechanistic studies. Development of precision biologics that specifically target FBs may be an effective strategy to avoid the side effects of broad cytokine blockade in the treatment of these inflammatory diseases of the dermis.

## Methods

### Study design

The aim of this study was to understand how the proinflammatory cytokine IL-17 can drive fundamentally different patterns of skin inflammation, as observed in HS versus psoriasis. To this end, we developed complementary murine systems of epidermal and dermal type 17 inflammation that reflect human disease inflammatory patterns. In vitro studies were used to understand the direct effect of IL-17 on different cell types. To demonstrate the relative importance of IL-17 recognition by different cell types, we developed conditional knockout mice lacking *Il17ra* in specific cellular compartments.

### Animals and animal care

All animal experiments were approved by the University of California, San Diego (UCSD) Institutional Animal Care and Use Committee. WT C57BL/6 mice were purchased from The Jackson Laboratory. *Krt14*-Cre, *Pdgfra*-Cre, *Adipoq*-Cre, and *Il17ra*-fl/fl mice on C57BL/6 background were purchased from The Jackson Laboratory and bred and maintained at UCSD. Mice were housed under a specific pathogen–free condition with 12hr light and 12hr dark cycle at 20–22°C and 30–70% humidity. Experimental and littermate control animals were age- and sex-matched 7–9-wk-old males and females. In some experiments, *Il17ra*-fl/fl mice (Cre^-^) were used in place of WT C57BL/6 mice.

### Mouse models

1-3 days prior to experiments, mice were randomly selected, and back skin was shaved, treated with depilatory cream (Nair), and rinsed with water. For the dermal type 17 inflammation model, 500 ng rmIL-17A (317-ILB-050; R&D) and/or 200 ng rmTNF (PMC3014; Thermo Fisher Scientific) was injected i.d. in 50μl PBS daily for three consecutive days. Skin was harvested 24hr after the third challenge for qPCR, flow cytometry, and immunofluorescence. For the epidermal type 17 inflammation model, 5% IMQ (51672414506; McKesson Medical Surgical) was applied topically on back skin daily for 5 days. Skin was harvested 24hr after the fifth challenge for qPCR, flow cytometry, and immunofluorescence. For *S. aureus* skin infection, 5e6 CFU mid-logarithmic growth phase strain USA300 was injected i.d. in 50μl PBS. Skin was harvested 48hr post-infection for immunofluorescence. For *C. acnes* skin infection, 1e7 CFU strain 61 (UCSD) was injected i.d. in 50μl PBS. 100μl of squalene was topically applied to the backs of mice 24hr before infection and every 24 hours thereafter. Skin was harvested 48hr post-infection for immunofluorescence. For *C. albicans* skin infection, 2e8 CFU strain 18804 (ATCC) was applied topically in 50μl PBS following stratum corneum removal sandpaper. Skin was harvested 72hr post-infection for immunofluorescence.

### RNA isolation, cDNA library preparation, and RT-qPCR

RNA from cells and tissues were preserved with RNAlater (AM7020; Thermo Fisher Scientific). Samples were lysed and RNA was isolated using PureLink Isolation kit (12183025; Thermo Fisher Scientific) and made into cDNA using Verso cDNA synthesis kit (AB1453/B; Thermo Fisher Scientific). qPCR was performed using the CFX96 Real-Time System (Bio-Rad) with SYBR Green Mix (QP1311; Biomiga). Housekeeping gene *Tbp* was used to normalize expression for mice. Specific primer sequences (IDT) are as follows:

*Lcn2*: (F: TGCCACTCCATCTTTCCTGTT; R: GGGAGTGCTGGCCAAATAAG)

*Cxcl1*: (F: CTGGGATTCACCTCAAGAACATC; R: CAGGGTCAAGGCAAGCCTC)

*Cxcl5*: (F: TGCCCTACGGTGGAAGTCATA; R: TGCATTCCGCTTAGCTTTCTTT)

*Cxcl12*: (F: TGCATCAGTGACGGTAAACCA; R: TTCTTCAGCCGTGCAACAATC)

*Saa3*: (F: TGCCATCATTCTTTGCATCTTGA; R: CCGTGAACTTCTGAACAGCCT)

*Il36a*: (F: GCAGCATCACCTTCGCTTAGA; R: CAGATATTGGCATGGGAGCAAG)

*Il36b*: (F: AGAGTATTCAAATGTGGGAACCG; R: GACCCATACCATCTGTTGTGAG)

*Il36g*: (F: TCCTGACTTTGGGGAGGTTTT; R: TCACGCTGACTGGGGTTACT)

*S100a8*: (F: AAATCACCATGCCCTCTACAAG; R: CCCACTTTTATCACCATCGCAA)

*S100a9*: (F: ATACTCTAGGAAGGAAGGACACC; R: TCCATGATGTCATTTATGAGGGC)

*Timp1:* (F: GCAACTCGGACCTGGTCATAA; R: CGGCCCGTGATGAGAAACT)

*Tnfrsf12a:* (F: GTGTTGGGATTCGGCTTGGT; R: GTCCATGCACTTGTCGAGGTC)

*Nfkbiz:* (F: GCTCCGACTCCTCCGATTTC; R: GAGTTCTTCACGCGAACACC)

*Zc3h12a:* (F: ACGAAGCCTGTCCAAGAATCC; R: TAGGGGCCTCTTTAGCCACA)

*C3:* (F: CCAGCTCCCCATTAGCTCTG; R: GCACTTGCCTCTTTAGGAAGTC)

### Histology and immunofluorescence

For H&E, samples were formalin-fixed paraffin-embedded (FFPE) and sectioned by the UCSD Tissue Technology Shared Resource Histology Core using standard protocols. Epidermal thickness was quantified using ImageJ. For immunofluorescence, skin biopsies were embedded fresh in OCT, frozen, and sectioned to 20μm at −20°C using a Leica CM1860 cryostat. Skin sections were briefly fixed in 4% paraformaldehyde, blocked with serum from secondary antibody host, stained with primary antibodies overnight at 4°C, secondary antibodies for 1h at room temperature, and nuclei were counterstained with DAPI. Epifluorescence images were taken using an EVOS5000. Brightness and contrast were adjusted slightly and applied equally across samples. Primary antibodies were as follows: LCN2 (1:100, AF1757; R&D), GR-1 (1:500, 108435, BioLegend), CXCL1 (1:100, PA586508, Thermo Fisher Scientific), Ly6G antibody (1:200, 127602, BioLegend). Secondary antibodies were as follows: Cy3 Donkey anti-Rabbit IgG (1:500, 406402; BioLegend), Donkey anti-Rat IgG (H+L) Highly Cross-Adsorbed Secondary Antibody, Alexa Fluor 488 (1:2000, A21208, Thermo Fisher Scientific), and Cy3 Goat anti-Rat IgG (1:500, 405408, BioLegend).

### scRNA-seq

FFPE skin biopsies were obtained as described above. Libraries were generated using the 10X Genomics Flex FFPE protocol (10X Genomics, Pleasanton, Calif) and subjected to 28 × 91 bp of sequencing according to the manufacturer’s protocol (Illumina NovaSeq [Illumina, San Diego, Calif]). Library preparation and next-generation sequencing were carried out in the Advanced Genomics Core at the University of Michigan. Reads were aligned to GRCm39 (refdata-gex-GRCm39-2024-A, available from 10X Genomics) using Cellranger multi software (v 9.0). Seurat 4.0 was used to merge and process the gene expression matrices. SoupX (v 1.6.2) was used to remove ambient RNA using default parameter^38^. ScDblFinder (v 1.16.0) was used to remove doublets, using default parameters^39^. Cells containing < 200 UMI and/or >25% mitochondrial genes were removed. The merged data matrices were then log-normalized, scaled, and principal component analysis was performed using 50 PCs^40^. Harmony (v 1.0.1) was used to perform batch effect correction, using donor as batch^41^. Clusters were labeled manually using known marker genes. Cell-cell communication analysis was performed using CellChat^13^. Pseudo-bulk differential expression analysis was performed using DESeq2^42^.

### RNA-Seq

Isolated RNA from cells in culture passing quality control (RNA integrity number > 8) underwent stranded mRNA sequencing on a Novaseq 6000 (Illumina) with paired-end 100 base-pair reads. RNA from mouse skin underwent Illumina Total RNA Prep (Ribodepletion). Reads were aligned to reference genome (mm10) using STAR^43^, and count tables were generated using FeatureCounts^44^. Data were normalized for gene length and sequencing depth by transcript per million (TPM) transformation, allowing gene-to-gene comparison and sample-to-sample comparison.

### Flow cytometry

Single-cell suspensions were prepared as described above. Cells were incubated with anti-CD16/CD32 (101302; BioLegend) for 10 min at 4°C to block non-specific Fc receptor binding and then incubated for 30 min with fluorochrome-conjugated antibodies diluted to 1:50 at 4°C including anti-Ly6G (127608; BioLegend), anti-CD11b (101212; BioLegend), and anti-CD45 (109828; BioLegend). Cell counting and flow cytometric acquisition was performed using a Novocyte (ACEA) flow cytometer. Data were analyzed using FlowJo (BD).

### Cell culture

All cells were grown in a humidified incubator at 5% CO2 and 37°C under sterile conditions. For primary fibroblast studies, murine neonatal (P1) cells were used. Primary dermal fibroblasts were isolated by our laboratory as previously described^45^ and used in passage 1. Cells were grown in Dulbecco’s Modified Eagle Medium (DMEM) supplemented with 10% FBS, Glutamax (35050061; Thermo Fisher Scientific), and antibiotic–antimycotic (15240062; Thermo Fisher Scientific). 2-day post-confluent cells were stimulated with recombinant cytokines and/or an adipogenesis-inducing cocktail. Adipogenesis was induced as described previously^46^. Following cytokine stimulation, cells were assayed at 24hr for qPCR and RNA-seq and CM harvested at 72hr for neutrophil migration. Recombinant cytokines used include rmIL-17A (50 ng/ml, 421-ML-100/CF; R&D) and rmTNF (20 ng/ml, PMC3014; Thermo Fisher Scientific).

### Neutrophil migration assay

Bone marrow was flushed from femurs and tibias of WT mice. Following red blood cell lysis, samples were filtered through a 40-μm strainer and resuspended at 2e6 cells/ml in DMEM with 0.5% FBS. 200 μl of the bone marrow suspension was loaded into the upper chamber of each well in a 3-μm 24-well transwell plate. 600 μl of fibroblast CM (0.5% FBS) was loaded into the bottom chamber. Cells were allowed to migrate for 3 h at 37°C. After 3 h, 60μl 0.5 M EDTA was added to lower chambers and plates were incubated at 4°C for 15 min. Migrated cells were collected from the lower chamber, counted, and stained for flow cytometric analysis as described above.

### Statistical analysis

Unless indicated otherwise, experiments were performed with at least biological triplicates and repeated at least twice. Statistical significance was calculated using R or GraphPad Prism with *P < 0.05, **P < 0.01, ***P < 0.001, and ****P < 0.0001. For qPCR and flow cytometry data in Figures 3, 4, and S2 fold-changes were calculated for each independent experiment prior to pooling. Analysis in Figure 1C-E analysis includes previously published data^8^ and data presented in Figure 3C-E and Figure 4C-E. Figure 2C analysis includes data presented in Figure 3I, Figure 4I, and Figure S2B,F.

### Study approval

All animal experiments were approved by the UCSD Institutional Animal Care and Use Committee (#S09074).

## Acknowledgments

We thank Professors Taylor Doherty and David Broide (UCSD) for flow cytometry resources. Bulk RNA-seq was supported by the UCSD Institute for Genomic Medicine Genomics Center and the NIH Shared Instrumentation Grant (#S10 OD026929). FFPE generation was supported by the UCSD Tissue Technology Shared Resource and the National Cancer Institute Cancer Center Support Grant (CCSG Grant P30CA23100). scRNA-seq was supported by the University of Michigan Advanced Genomics Core, the University of Michigan Single-Cell Spatial Analysis Program, and the National Cancer Institute (P30CA046592) using the Cancer Center’s Single-Cell and Spatial Analysis Shared Resource. K.J.C. is supported by NIH T32AI007036 and the Hartwell Foundation, and R.L.G. is supported by NIH R01AI153185, R37AI052453, U01AI152038, and P50AR080594. J.E.G, L.C.T, and J.M.K are supported by NIH-P30-AR075043. Figure models were created using https://BioRender.com.

## Author contributions

K.J. Cavagnero—conceptualization, investigation, resources, formal analysis, visualization, writing (original draft), writing (review & editing); J.M. Kahlenberg, C. Aguilera—resources; J. Fox, J. Kirma, H. Jo, F. Li—investigation; R. Bogel—resources, formal analysis. L.C. Tsoi, J. Gudjonsson—supervision, resources. R.L. Gallo—supervision, conceptualization, and writing (review & editing).

## Data availability

Novel genomic data presented here will be made available in the National Center for Biotechnology Information Gene Expression Omnibus database following publication.

