## Supplemental Figures for "Disease context dictates the cellular targets of IL-17 in inflammatory skin disease"

### Fig. S1

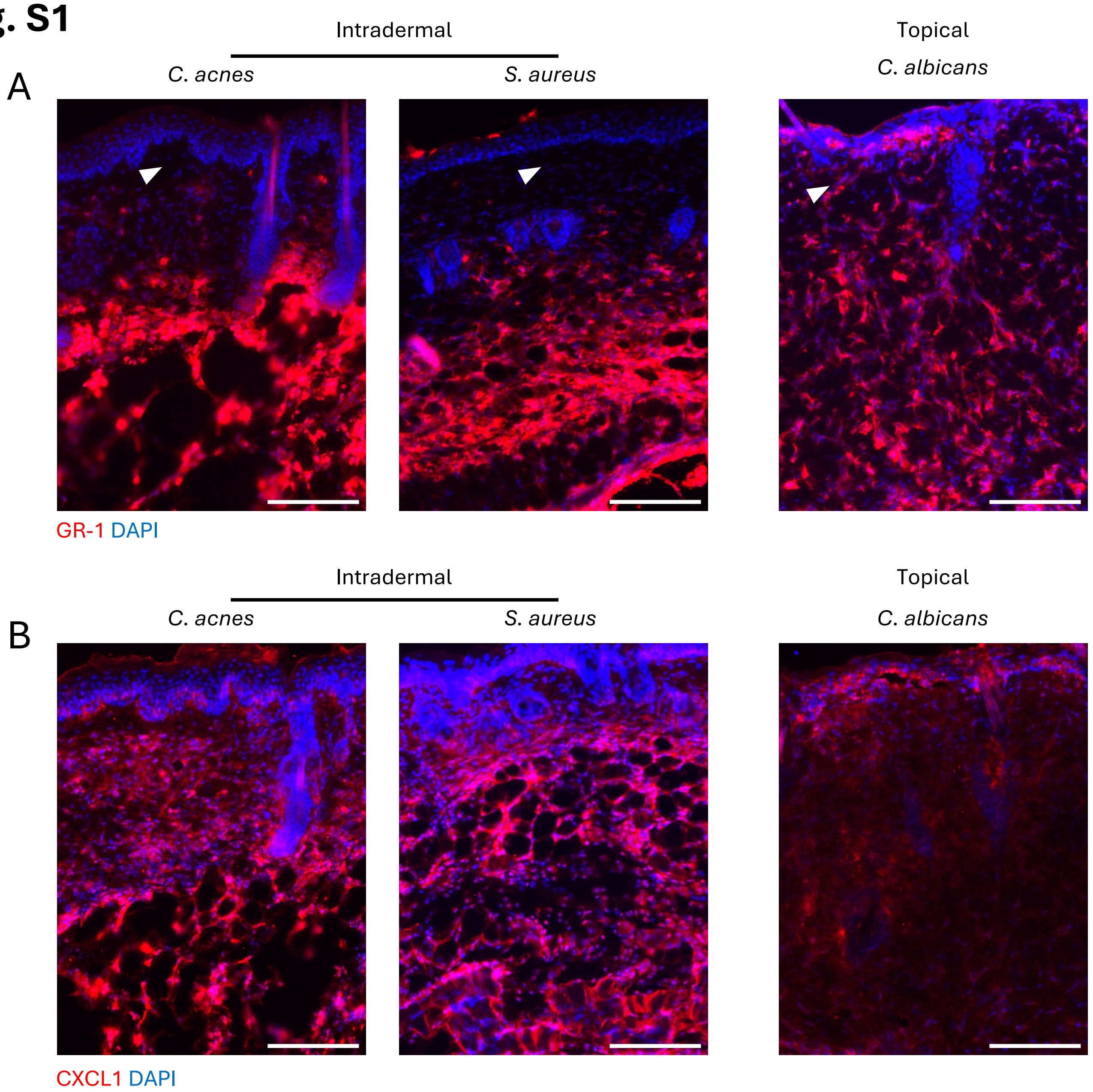


**Fig. S1 | Infection depth dictates dermal versus epidermal inflammatory patterns.** Representative mouse skin immunostaining for GR-1 (A) and CXCL1 (B) following infection. Left = intradermal challenges. Right = epi-cutaneous (topical) challenge. Scale bars, 150um. Arrows denote region with marked difference in sub-epidermal inflammation.

Fig. S2

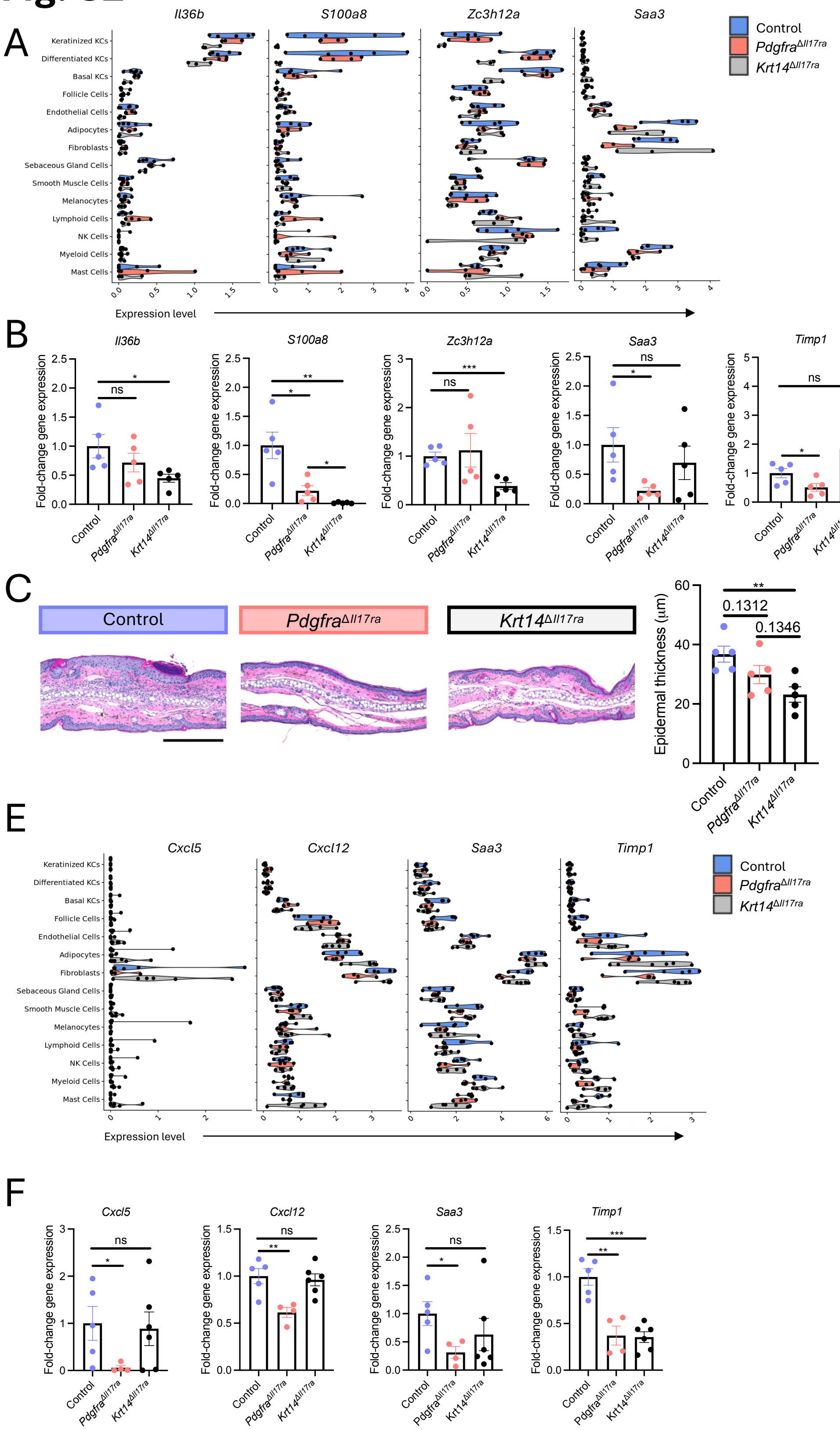


**Fig. S2 | Role of dermal fibroblasts versus keratinocytes in dermal and epidermal type 17 inflammation.** (A-B) Topical IMQ experimental system (dorsal skin). (A) scRNA-seq pseudo-bulk gene expression. Each point represents 1 mouse. (B) Skin qPCR gene expression. (C-D) Topical IMQ experimental system (ear skin). (C) Representative skin H&E staining (left) and epidermal thickness quantification (right). (D) Skin qPCR gene expression. (E-F) i.d. IL-17/TNF experimental system (dorsal skin). (E) scRNA-seq pseudo-bulk gene expression. Each point represents 1 mouse. (F) Control = $Cre^-$ littermate. ns (not significant), $*p < 0.05$, and $**P < 0.01$ using unpaired t-test.

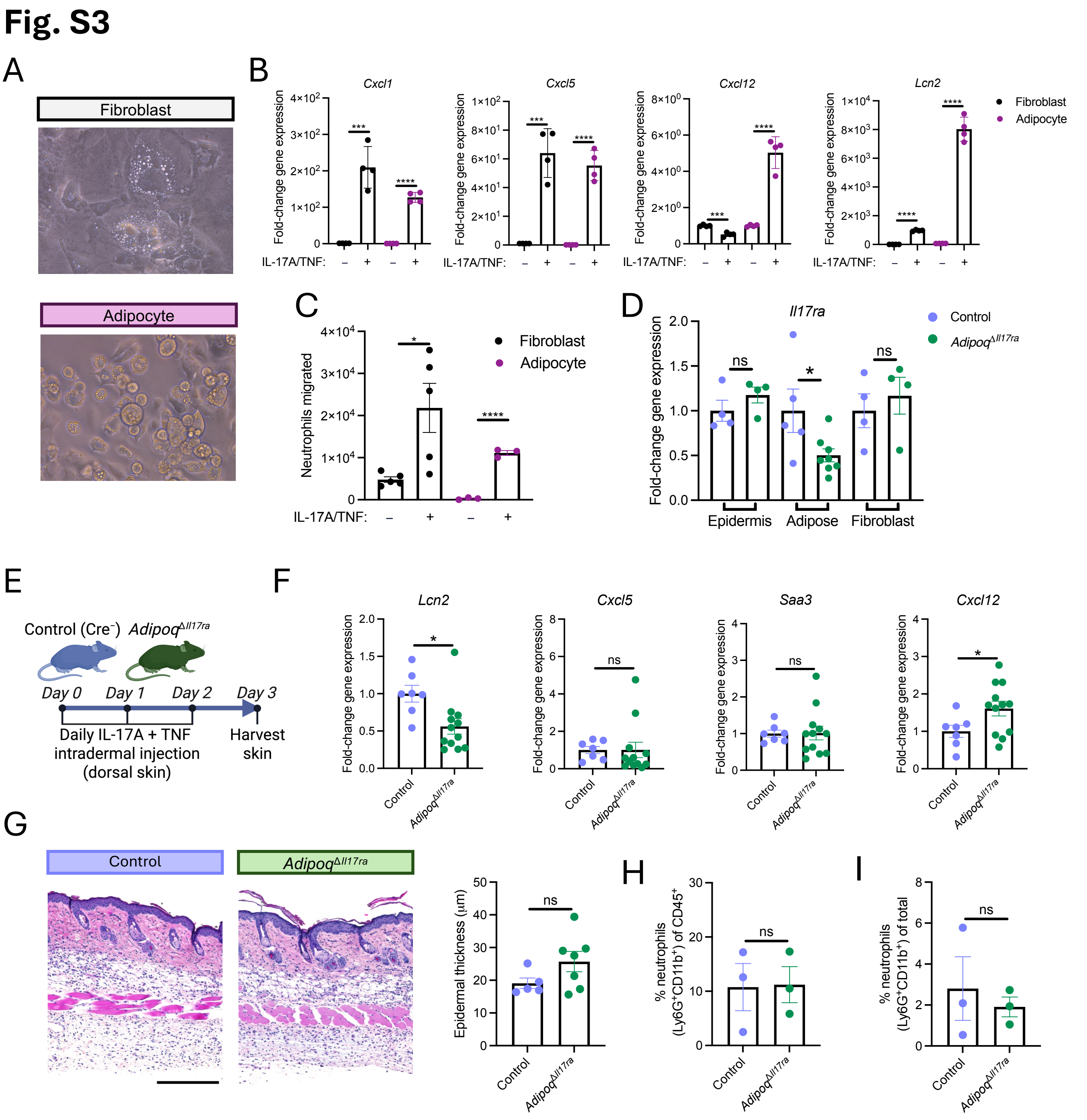


**Fig. S3 | Role of dermal adipocytes in dermal type 17 inflammation.** (A) Undifferentiated and adipocyte-differentiated primary mouse dermal FB. (B) qPCR gene expression following treatment with IL-17/TNF. (C) Neutrophil transwell migration assay using conditioned media from fibroblasts or adipocytes that were treated with IL-17 and TNF. (D) qPCR gene expression using isolated skin compartments from naïve mice. (E-I) i.d. IL-17/TNF experimental system. (E) Experimental schematic.

(F) Skin qPCR gene expression. (G) Representative H&E (left) and epidermal thickness quantification (right). Scale bar, 300um. Flow cytometric quantification of the frequency of neutrophils (Ly6G+CD11b+) of live CD45+ skin cells (H) and total neutrophils per 1.5cm2 skin biopsy (I). Control = $Cre^{-}$ littermate. ns (not significant), *p < 0.05, ***P < 0.001, and ****P < 0.0001 using unpaired t-test. i.d., intradermal.
